# Transcription repressor protein ZBTB25 interacts with HDAC1 in macrophages infected with *Mycobacterium tuberculosis,* and its inhibition leads to autophagy and killing of the intracellular pathogen

**DOI:** 10.1101/2020.06.22.164350

**Authors:** Aravind Madhavan, K B Arun, Akhil Raj Pushparajan, M Balaji, R Ajay Kumar

**Author notes:** authors contributed equally (Aravind Madhavan, K B Arun). corresponding author R. Ajay Kumar, Mycobacterium Research Laboratory, Rajiv Gandhi Centre for Biotechnology, Thiruvananthapuram – 695014, Kerala, India.

## Abstract

Downregulation of host gene expression is one of the key strategies adopted by intracellular pathogens such as *Mycobacterium tuberculosis* (MTB) for their survival and subsequent pathogenesis. In a previous study, we have shown that HDAC1 levels go up in macrophages infected with MTB and it hypoacetylates histone H3 at the promoter of *IL-12B* gene leading to its downregulation. Here we show that after infection with MTB, the levels of the phosphorylated form of HDAC1 increase significantly in macrophages. Employing immunoprecipitation and LC-MS/MS, we found transcriptional repressor protein ZBTB25 associates with HDAC1 silencing complex along with transcriptional corepressor Sin3a. By chromatin immunoprecipitation and PCR analyses, we found that phosphorylated HDAC1, Sin3a, and ZBTB25 are recruited to the promoter of *IL-12B* to downregulate its expression in infected macrophages. Knocking down of *ZBTB25* enhanced release of IL-12p40 from infected macrophages. Interestingly, the treatment of infected macrophages with CI994 (inhibitor of HDAC1) or dithiopyridine (inhibitor of ZBTB25) promoted the colocalization of LC3 (microtubule-associated protein 1A/1B-light chain 3, a marker for autophagy) and MTB in autophagosomes. Induction of autophagy resulted in the killing of intracellular MTB. Enhanced phosphorylation of JAK2 and STAT4 was observed in macrophages upon CI994 and dithiopyridine treatment, and inhibition of JAK2/STAT4 negated the killing of intracellular mycobacteria suggesting a possible role of these proteins in the autophagy-mediated killing of intracellular MTB.

## Introduction

Tuberculosis (TB) caused by *Mycobacterium tuberculosis* (MTB) is a prominent cause of death worldwide; especially in tropical regions. Although effective anti-TB drugs and BCG vaccine have been in use for almost a century, TB still remains a global malady that cannot be eradicated. The current therapeutic drugs inhibit MTB by interfering with critical pathways such as DNA metabolism and mycolic acid biosynthesis. Typically treatment of TB involves administration of multiple drugs over a period of 4-6 months. This, combined with side effects of the drugs, leads to noncompliance which is one major reason for emergence of drug resistant TB. The ability of MTB to remain within the host in a dormant state for a long time, and to get reactivated when the body’s defences are compromised, is another reason for the success of MTB as a recalcitrant pathogen. Novel and fast acting anti-TB molecules against metabolically active and dormant bacteria are required for effective control of TB. In recent times, however, an alternative approach namely host-directed therapy has emerged as a promising treatment method which, unlike the current anti-TB drugs, targets host factors which facilitate the survival and pathogenicity of MTB (Zumla et al, 2016; Kaufman et al, 2018).

After invading the host, MTB inhabits membrane-bound phagosomes and employs mechanisms to evade the innate immune responses that trigger macrophages (Deretic & Levine, 2009; Barry et al, 2010). After phagocytosis, the primary host defense mechanism is initiated by the secretion of pro-inflammatory cytokines TNF, IL-1β, IL-6, and IL-12. Pro-inflammatory cytokines play a key role in initiating antimicrobial responses (Jayaraman et al, 2013; Law et al, 1996; Robinson et al, 2008). The T-cell-mediated immunity augments the capability of macrophages to clear bacteria by activating Th1 responses and boosting its cytotoxic activity (Salgame et al, 2005).

Recent studies reveal that intracellular pathogens suppress host immune functions by targeting the host chromatin and epigenetic regulators (Kathirvel and Mahadevan, 2016). In a previous study we had shown that infection of human macrophages with virulent MTB causes the levels of HDAC1 to increase significantly and repress expression of *IL-12B* gene (Chandran et al., 2015). HDAC1 has been shown by numerous studies to be the catalytic core of many co-repressor complexes involved in the downregulation of eukaryotic gene expression.

Microbial pathogens have developed several strategies to take control of the host immune system. Suppression of host gene expression by HDACs plays a significant role in keeping the host immune system in check. Epstein-Barr virus proteins TRF2 (Zhou et al, 2009), and EBNA3C (Knight et al, 2003) bind to HDAC1 complex and these interactions are necessary for viral replication in host and infection. Terhune et al (2010) identified cellular proteins RBBP4, CHD4 and Human Cytomegalo Virus (HCMV) protein pUL29/28/38 interact with HDAC1 upon HCMV infection to stimulate the accumulation of immediate-early viral RNAs. 2-Aminoacetophenone, a quorum sensing molecule from *Pseudomonas aeruginosa* regulate host HDAC1 expression and enhances the interaction between cellular p50 protein and HDAC1 leading to repression of the proinflammatory responses (Bandyopadhaya et al, 2017). Ankyrin molecule from *Anaplasma phagocytophilum* induces the recruitment of HDAC1 complex to promoters of antimicrobial defense genes (Rennoll-Bankert et al, 2015). Our own study has shown that HDAC1 is recruited to the promoter of *IL-12B* to downregulate its expression in MTB-infected macrophages (Chandran et al, 2015). However, it remains to be seen if MTB proteins are associated with the complex in infected cells. Also, the identity of host proteins associated with HDAC1 during MTB infection and their role in the progress of infection/pathogenesis are also not known. Although our study has shown that knocking down or inhibiting HDAC1 resulted in the enhanced killing of intracellular bacteria, we propose that inhibition of HDAC1-associated proteins, rather than inhibiting HDAC1 itself which has a broader role in cellular gene regulation, maybe a more specific strategy for host-directed anti-TB therapy.

The present study, in which we analyzed the HDAC1 associated proteins in MTB-infected macrophage cells, reveals that ZBTB25, a repressive transcription factor associates with HDAC1 repressor complex. ZBTB25 is a member of the broad complex tram track bric-a-brac/poxvirus and zinc finger (BTB/POZ) transcription family, and these proteins possess a DNA binding zinc finger motif at the C-terminal and a protein binding BTB/POZ domain at the N-terminal. The zinc finger identifies and binds to the specific DNA sequences, while the BTB/POZ domain helps in the homodimerization and/or heterodimerization and interaction with other proteins (Chang et al, 1996). The human genome encodes about 60 genes under the ZBTB family (Beaulieu and Sant’Angelo, 2011) although the function of most of the members is yet to be ascertained. In the present study, we show that ZBTB25, along with HDAC1, binds to *IL-12B* gene promoter and represses its expression. Inhibition of ZBTB25 derepresses the expression of *IL-12B* gene and also activates autophagy-associated genes. Because of its dual role in HDAC1-mediated repression of genes and autophagy, we propose that ZBTB25 may be a potential therapeutic target for TB treatment.

## Results

### HDAC1 in macrophages is phosphorylated upon MTB infection

Enzymatic activity and repressor complex formation of HDAC1 are controlled by phosphorylation. To delineate if HDAC1 is phosphorylated during MTB infection, human macrophages were infected with MTB H37Rv, and a time-course analysis of phosphorylated HDAC1 (pHDAC1) was carried out from 0 to 48 h by western blot analysis of uninfected and MTB-infected macrophages. At 24 h post-infection (PI), the level of pHDAC1 increased by 4-fold in macrophages infected with virulent MTB (Fig.1a & 1b).

**Figure 1.**
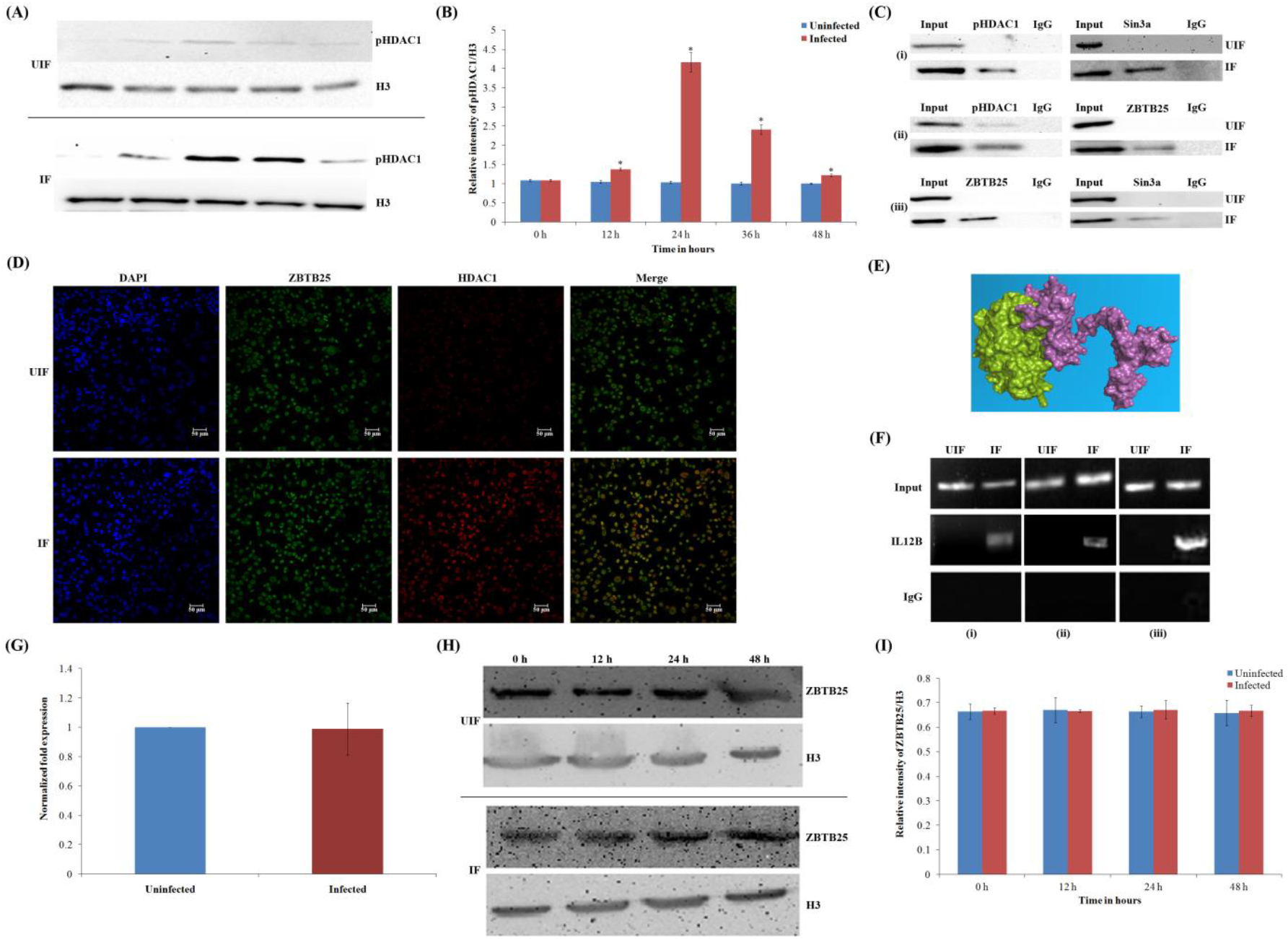
Phosphorylated HDAC1 interacts with ZBTB25 and they are recruited to *IL-12B* promoter. (A) Time course of levels ofphosphorylated HDAC1 in MTB-infected macrophages. pHDAC1 expressionin uninfected (UIF) and infected (IF) samples;(B) Densitometric analysis of pHDAC1 bands normalized with that of histone H3. Each value represents mean ± SD from triplicate measurements.*Significantly different from UIF sample, p≤0.05; (C) Co-immunoprecipitation analysis of the association between ZBTB25 and SIN3A-HDAC1 complex. Whole-cell lysates were immunoprecipitated with the antibodies against (i) ZBTB25, (ii) Sin3a;(iii) HDAC1.IgGwas used as the negative control. Immunocomplexes were then probed with antibodies as indicated; (D) Immuno-cytochemical imaging shows HDAC1 co-localizes with ZBTB25 inside the nucleus of macrophages infected with MTB.The cells were visualized by confocal microscopy at 24 h PI; (E) Docking analysis shows that HDAC1 can interact with ZBTB25;(F) Status of ZBTB25, HDAC1 and Sin3a recruitment on the *IL12B* promoter by ChIP. ChIP with (i) ZBTB25, (ii) HDAC1, and (iii) Sin3a; (G)Status of expression of *ZBTB25* in macrophages upon MTB infection; (H) Real time qPCR of ZBTB25 and western blot; (I)Densitometric analysis of ZBTB25 bands normalized with that of histone H3. Each value represents mean ± SD from triplicate measurements.

To find out whether it is the pHDAC1 that is recruited to *IL-12B* promoter in macrophages during MTB infection, we performed chromatin immunoprecipitation (ChIP) with antibodies against pHDAC1 followed by a PCR for the promoter sequence. Amplification of the *IL-12B* promoter region confirmed the recruitment of pHDAC1 to it (Fig.1f (II)).

### Phosphorylated HDAC1 interacts with ZBTB25, and both are recruited to *IL-12B* promoter

To identify the interacting partners of HDAC1 during MTB infection, we infected THP1-derived macrophages with MTB H37Rv and immunoprecipitated pHDAC1-associated proteins from cell extracts with pHDAC1-specific polyclonal antibodies. Subsequent LC-MS/MS analysis detected and identified HDAC1 and two other macrophage proteins namely ZBTB25 and Sin3a (Supplementary Table 1) in the precipitate. Interaction of ZBTB25 with HDAC1 and Sin3a was confirmed by reciprocal co-immunoprecipitations (Fig.1c). To further investigate the association of HDAC1 with ZBTB25, we carried out an immunofluorescence microscopy analysis. Confocal images of the macrophages infected with MTB showed that ZBTB25 and HDAC1 colocalize within the nucleus at 24 h PI (Fig. 1d). Such colocalization was not observed in uninfected cells. Interaction of the HDAC1 and ZBTB25 was analysed employing in silico docking analysis which showed that HDAC1 and ZBTBT25 interact with each other strongly with a docking score of −914.5 Kcal/mol (Fig.1e and Supplementary Fig. 1. Following infection, ZBTB25 and Sin3a were found to be recruited to the *IL-12B* promoter at 24 h PI. This was confirmed by chromatin immunoprecipitation (ChIP) of macrophage DNA with antibodies against ZBTB25 and Sin3a, followed by a PCR for these promoter sequences (Fig.1f). The levels of ZBTB25 in the uninfected and infected macrophages were evaluated using real-time PCR, and a time-course analysis was done using western blotting with ZBTB25 specific antibodies (Fig.1g). We found no significant difference in the levels of ZBTB25 in MTB infected and uninfected macrophages (Fig.1h & 1i).

### Knocking down of ZBTB25 restores the expression of *IL-12B* and reduces the intracellular survival of MTB

Since we observed an interaction between HDAC1 and ZBTB25, we tested whether ZBTB25 plays a role in the survival of MTB inside the macrophages. To investigate this, we knocked down *ZBTB25* in THP1-derived macrophages and infected them with virulent MTB. The efficiency of knockdown was confirmed by quantitative reverse transcription-PCR (qRT-PCR), western blotting, and confocal microscopy (Fig. 2a, 2b, and 2c). We observed that ZBTB25, HDAC1, and Sin3a were not recruited to the promoter of *IL-12B* in macrophages in which ZBTB25 was knocked down (Fig. 2d). As expected, knockdown of *ZBTB25* significantly enhanced *IL-12B* mRNA and IL-12p40 protein levels in infected cells as confirmed by qRT-PCR (Fig.2e) and ELISA (Fig. 2f). Colony-forming units (CFU) of intracellular MTB were enumerated by lysing the infected macrophages at 24 h PI and plating them on Middlebrook 7H10 agar. Macrophages treated with scrambled siRNA and infected with MTB served as control. A significant decrease was observed in the number of CFU in macrophages in which *ZBTB25* was knocked down (Fig. 2g).

**Figure 2:**
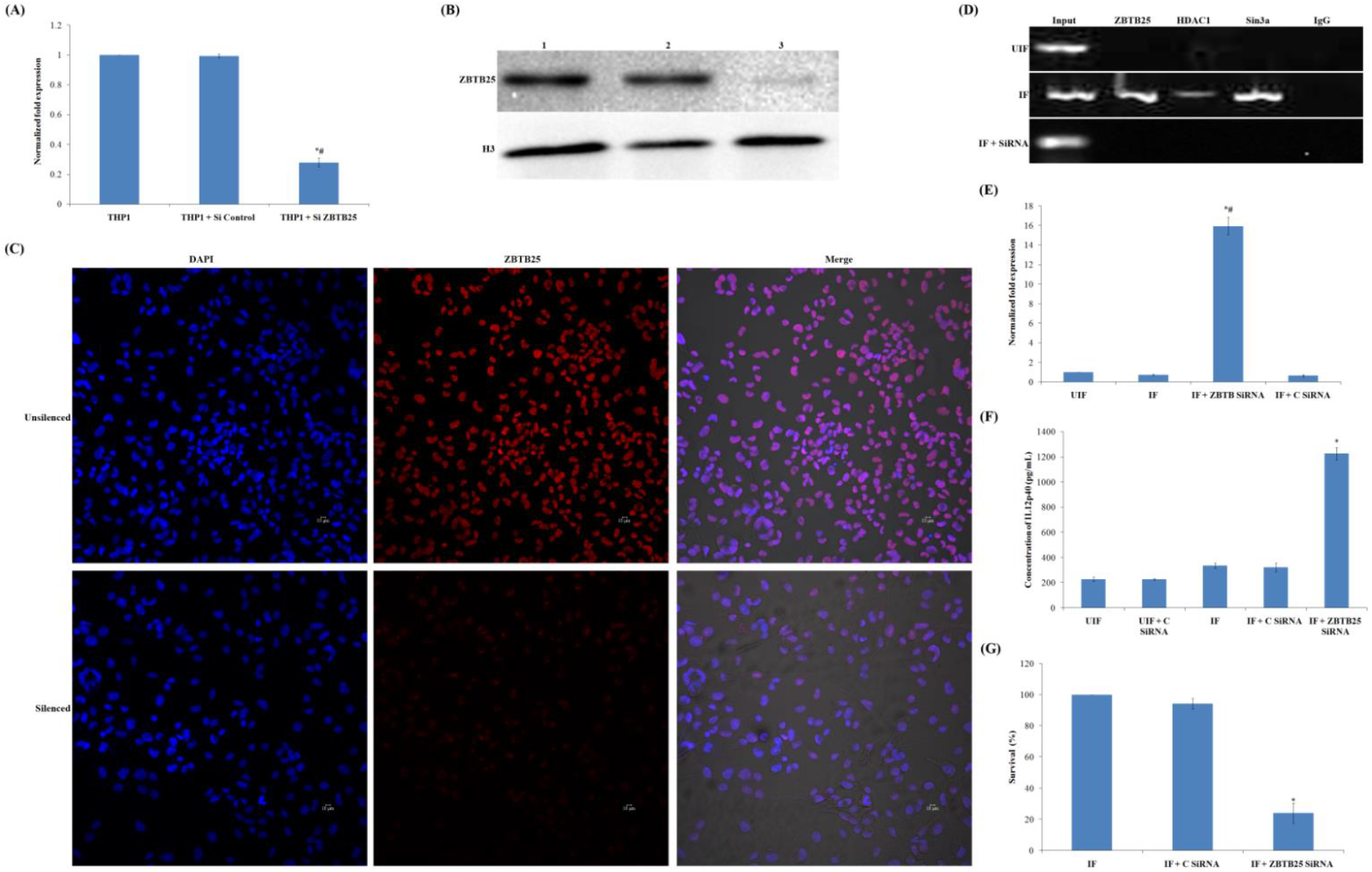
Knocking down of ZBTB25 represses *IL-12B* and reduces the intracellular survival of MTB. **(A)** THP1 macrophage cells were transfected with *ZBTB25* SiRNA or scrambled SiRNA control using Hiperfect transfection reagent.Efficiency of knock down was confirmed by qPCR.The data are representative of three independent experiments with similar results. Expression of *ZBTB25* in macrophages in which it is knocked down is significantly less from the controls. (B) Western blot analysis of macrophage cells in which the expression of endogenous ZBTB25 was knocked down using ZBTB25 SiRNA. Lane 1: Normal macrophages, Lane 2: Macrophages transfected with scrambled SiRNA control, and Lane 3: Macrophages transfected with ZBTB25 SiRNA. (C) Immuno-cytochemical staining with anti-ZBTB25 antibody and DAPI of control THP1 macrophages, and cells in which ZBTB25 was knocked down showed reduced fluorescence compared to the control THP1 macrophages (D) knocking down of ZBTB25 blocks the recruitment of the HDAC1 silencing complex to the*IL-12B* promoter. Status of (i) ZBTB25, (ii) HDAC1, and (iii) Sin3a on the *IL-12B* promoter by ChIP followed by PCR.(E) *IL-12B* expression is upregulated in MTB-infected macrophages upon *ZBTB25* inhibition. qPCR of *IL-12B* when *ZBTB25* is knocked down.*#*IL-12B* expression is significantly different from uninfected (UIF) and infected (IF) samples, p≤0.05. (F) ELISA of IL-12p40 when *ZBTB25* is knocked down in macrophages. *IL-12p40 expression is significantly different from infected (IF) sample, p≤0.05. (G)Survival of MTB decreases in macrophages when ZBTB25 is knocked down. Macrophages in whichZBTB25 wasknocked down were infected with MTB. Macrophages not treated with siRNA but infected with MTB were also kept as control.Intracellular bacterial viability was determined based on the number of CFUs. Data shown here are the mean ± SD of three independent experiments. *Survival is significantly different from both controls, p≤0.05.

### Treatment of infected macrophages with dithiopyridine and CI994 enhances intracellular clearance of MTB

To test the effect of inhibition of ZBTB25 in the survival of MTB in macrophages, infected macrophages were treated with dithiopyridine, a zinc ejector drug that blocks zinc finger domain, after phagocytosis. Intracellular growth was significantly reduced (35%) when the cells were treated with 20 μM dithiopyridine at 24 h PI compared to the MTB- infected but untreated macrophages (Fig.3a). In addition we evaluated the effect of CI994 an HDAC1 inhibitor, on intracellular survival of MTB. At 24 h PI macrophages that were treated with CI994 showed a significant decrease (47%) in the number of intracellular MTB (Fig.3a). Interestingly treatment with a combination of dithiopyridine and CI994 showed a synergistic effect on the expression of *IL-12B*, IL12p40 levels, and the intracellular MTB survival (Figs. 3a, 3b, and 3C, respectively). The CFU count indicated a significant reduction in the number (28% survival) of MTB inside the macrophages compared to the macrophages treated with dithiopyridine and CI994 individually (Fig.3a). In the presence of dithiopyridine and CI994, recruitment of HDAC1 complex proteins (HDAC1, ZBTB25 and Sin3a) to the *IL-12B* promoter was not observed (Fig.3d).

**Figure 3.**
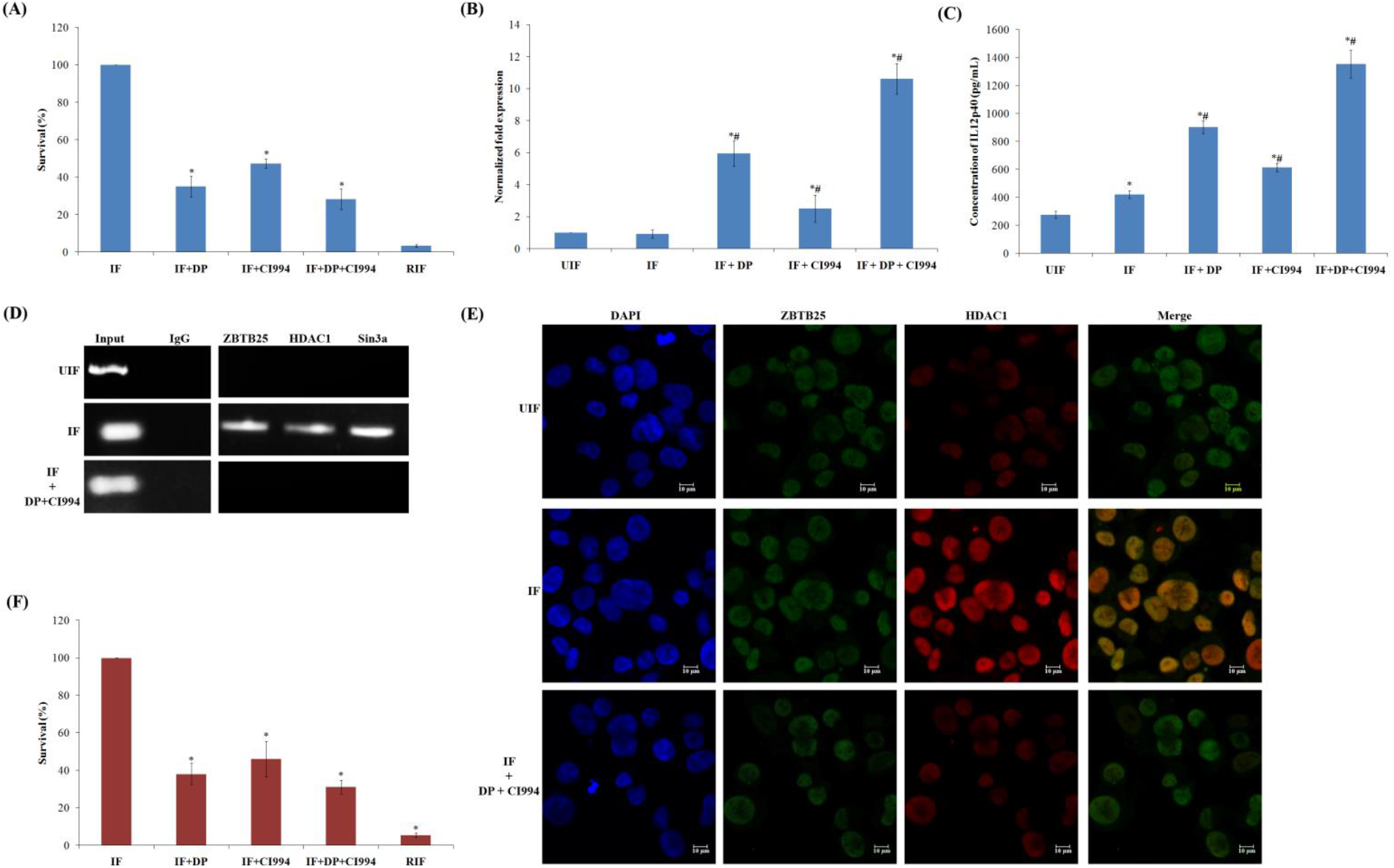
Treatment of infected macrophages with dithiopyridine (ZBTB25 inhibitor) and CI994 (HDAC1 inhibitor) enhanced the intracellular clearance of MTB. THP1-derived macrophages were infected with the virulent strain of MTB H37Rv. After MTB infection, cells were treated with dithiopyridine (DP) and/or CI994 orrifampicinfor 24 h. (A) Intracellular bacterial viability was determined based on the number of CFUs.Isolation of bacilli from the macrophages was carried out at 24 h. The number of viable bacilli in each of the plate was assayed by plating lysed macrophages on 7H10 agar plates and incubating the plates at 37°C and counting the CFUs. Values are shown from 3 independent experiments (mean ± SD). * Survival is significantly different from infected samples (IF), p≤0.05;(B) qPCR for the expression of the *IL-12B* mRNA transcript (normalized to β-actin mRNA expression).*, #*IL-12B* expression is significantly different from uninfected (UIF) and infected samples (IF) respectively, p≤0.05;(C) ELISA of IL-12p40 after inhibitor treatment. *, # IL-12p40 expression issignificantly different from uninfected (UIF) and infected samples (IF) respectively, p≤0.05; (D) DP/CI994 treatment blocks the recruitment of the HDAC1 silencing complex to *IL-12B* promoter. Status of (i) ZBTB25, (ii) HDAC1, and (iii) Sin3a on the *IL-12B* promoter by ChIP PCR; (E) Immuno-cytochemical imaging shows HDAC1 is not colocalizing with ZBTB25 inside the macrophage nucleus after DP and CI994 treatment; (F) Intracellular bacterial viability in MTB infected PBMC after treatment with DP,CI994 and rifampicin was determined based on the number of CFUs. * Survival is significantly different from infected sample (IF), p≤0.05.

Treatment with inhibitors disrupted the interaction and colocalization of ZBTB25 and HDAC1 in the nucleus, which was confirmed by confocal microscopy (Fig.3e). Toxicity of dithiopyridine towards THP-1 cells was tested by MTT assay, and that towards MTB H37Rv was tested by resazurin microtitre assay (REMA) and confirmed that the drug concentration used was not toxic to THP1-derived macrophages and MTB (Supplementary Fig. 2, 3, 4 and 5).

Consistent with our observation in THP1-derived macrophages, we found a similar trend when peripheral blood mononuclear cells (PBMC) were infected with MTB and treated with these inhibitors. Treatment with dithiopyridine and CI994 resulted in the killing of 62% and 54% of MTB, respectively (Fig.3f), and the combination of dithiopyridine and CI994 enhanced the killing of intracellular MTB to 70% in PBMC. The levels of IL-12p40 were found to be significantly high in MTB-infected macrophages treated with dithiopyridine and CI994, compared to those in infected but untreated cells (Supplementary fig. 6).

### Autophagy is activated when *IL-12B* expression is restored in macrophages

Treatment of macrophage with dithiopyridine (20 μM) and CI994 (15 μM) significantly enhanced *IL-12B* expression and decreased the survival of MTB. Since dithiopyridine and CI994 are found to increase the IL-12p40 levels, and since it is reported that IL-12 mediates killing of intracellular mycobacteria through autophagy, we investigated if dithiopyridine and CI994 can induce autophagy in THP-1-derived macrophage cells. Autophagosome formation is regulated by two critical proteins, namely ATG5 and Beclin 1. Expression of *BECN1* (encoding Beclin 1) and *ATG5,* and their product levels (Fig. 4a and 4b) were low in macrophages infected with MTB. However, when they were treated with dithiopyridine alone or in combination with CI994, expression of both the genes, and their protein levels, were significantly elevated (Fig.4a and 4b; Fig. 4c and 4d). Thus, treatment with dithiopyridine and CI994 was able to overcome the MTB-mediated suppression of the autophagy genes *ATG5* and *BECN1* in human macrophages. Confocal microscopy revealed that the levels of BECN1 increased after the drug treatment (Fig.4e). LC3 is another key protein required for autophagosome formation and is considered an autophagy marker. MTB-infected macrophages showed reduced levels of LC3 (Fig. 5a). Interestingly dithiopyridine or CI994 treated cells could overcome this downregulation, and LC3 was found to be localized as punctate structures in the cells, and the number of punctae was more in cells treated with both the drugs (Fig. 5a).

**Figure 4.**
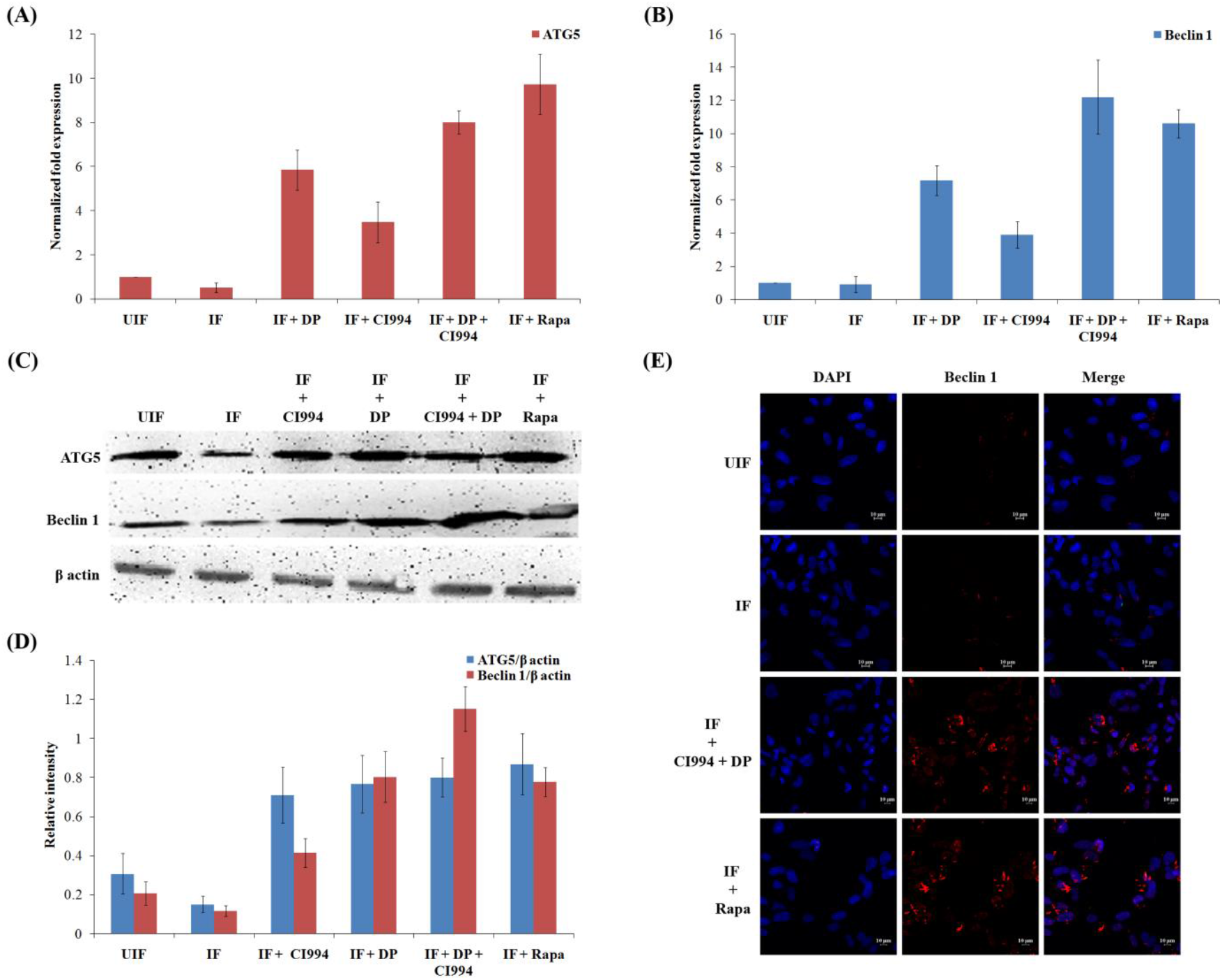
Dithiopyridine and CI994induce autophagy in human macrophages. (A) Dithiopyridine/CI994 treatment of infected macrophagesupregulated the expression of the autophagy-related genes Beclin 1 and ATG5. Differentiated human THP1 cells were infected with the virulent strain of MTB H37Rv for 4 h and treated with Dithiopyridine and/CI994 or rapamycin (Rap). The mRNA levels of autophagy-related genes ATG5 and (B) Beclin1 (normalized to β-actin expression) in THP1-derived macrophages were measured by qPCR. Results are shown from 3 independent experiments (mean ± SD); (C) Western blot of ATG5,Beclin 1 and β-actin; (D) Densitometric analysis of ATG5 and Beclin 1 bands normalized with that of β-actin. Each value represents mean ± SD from triplicate measurements;(E) Immuno-cytochemical imaging shows expression of Beclin 1 in variously infected macrophages at 24 h PI.

**Figure 5.**
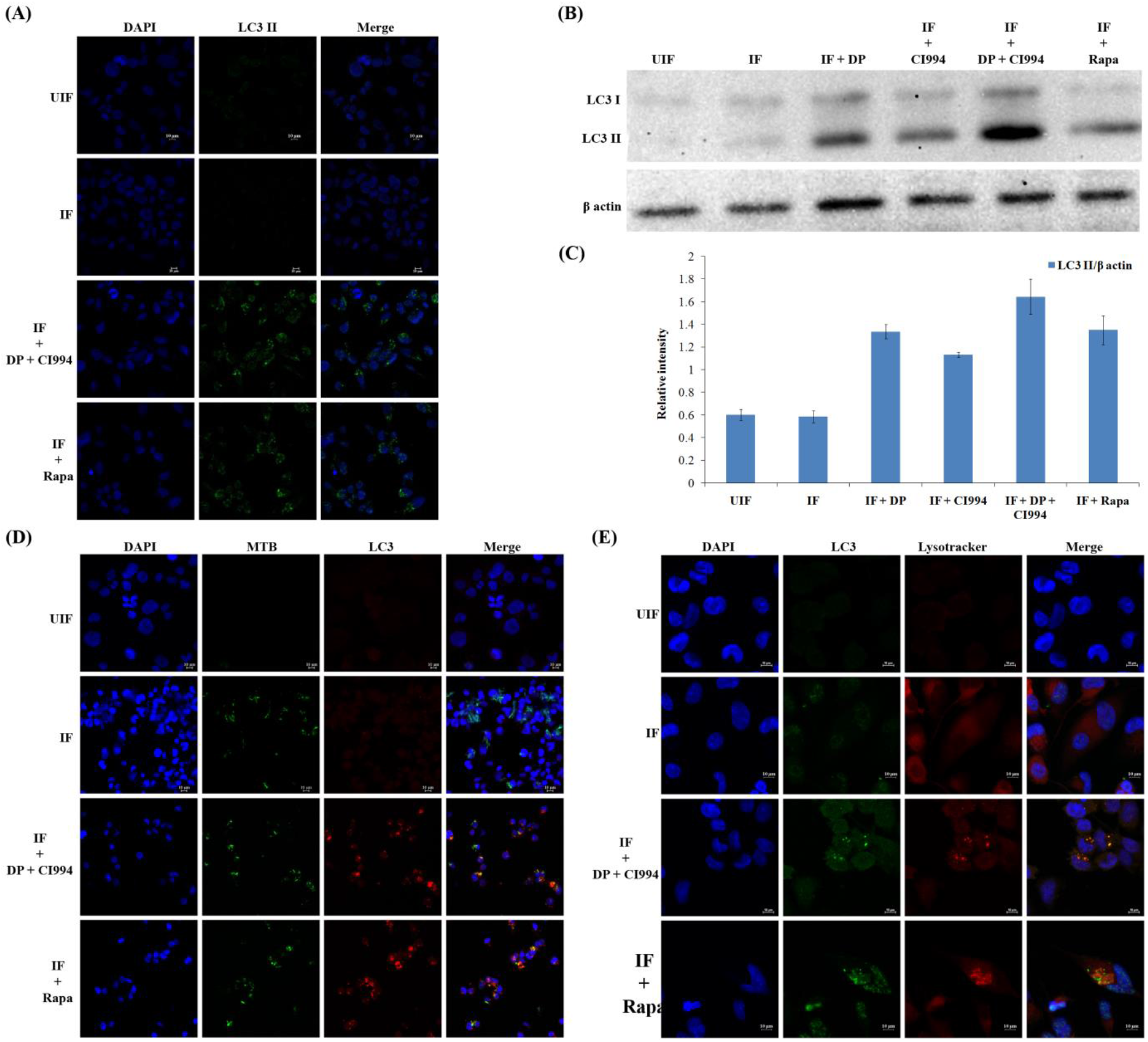
Dithiopyridine/CI994-induces autophagicclearacnce of bacteria by enhancing the expression of LC3 and MTB-LC3 co-localization in human macrophages. THP1-derived macrophages were infected with the virulent strain of MTB expressing GFP for 4 h and treated with dithiopyridine (DP), CI994 and rapamycin (Rapa). (A) The cells were fixed and stained with DAPI to visualize the nuclei (blue), and with anti-LC3 followed by the addition of FITC conjugated Mouse IgG (green). One representative immunofluorescence image out of 3 independent replicates is shown; (B) A representative western blot shows the conversion of LC3-I to LC3-II, and β-actin from 3 independent experiments; (C) Densitometric analysis of LC3-II bands normalized with that of β-actin.Each value represents mean ± SD from triplicate measurements; (D) Distribution of colocalization of GFP expressing MTB and LC3 (red) in macrophages at 24 h PI. (E). Macrophages were immunostained for LC3 (green) and LysoTracker-red to detect LC3 and lysosome-containing cells.

LC3 is usually located in the cytoplasm of cells as LC3-I, which is a non-lipidated form. After the induction of autophagy, LC3-II is formed by the covalent binding of phosphatidylethanolamine to LC3 (Runwal et al, 2019). This lipidated form of LC3 is associated with the membrane of the autophagosome. We analyzed the LC3-I to LC3-II conversion by western blot analysis (Fig.5b and 5c). MTB infected macrophages treated with dithiopyridine and CI994 enhanced the formation of LC3-II. Subsequently, employing fluorescence microscopy, we tracked the association of GFP-expressing MTB with autophagosomes in infected macrophages upon treatment with the inhibitors. Dithiopyridine and CI994 treatment caused an increase in the number of cells with LC3 punctae, and showed an increase in the colocalization of GFP-expressing MTB with LC3 autophagosomes (Fig. 5d) compared to untreated cells. As autophagy progresses, autophagosomes combine with lysosomes to form autolysosomes that carry LC3 and various lysosome-associated components. To find whether dithiopyridine/CI994 treatment and IL-12p40 restoration affect autolysosome formation, we analyzed the colocalization of endogenous LC3 and lysosomes by immunostaining with LysoTracker followed by fluorescence microscopy. The images (Fig. 5e) clearly show that the LC3 vesicles colocalized with lysosomes.

### IL-12p40-mediated activation of autophagy depends on JAK-STAT pathway

To find the role of pJAK2 and pSTAT4 in the activation of IL-12p40-mediated induction of autophagy, we assayed their levels in macrophage upon MTB infection by western blotting (Fig. 6a, 6b and 6C). We observed a significant increase in the levels of phosphorylation of pJAK2 and pSTAT4 upon treatment of infected macrophages with dithiopyridine and CI994, which suggests the involvement of IL-12 in the phosphorylation of JAK2 and STAT4. Treatment of infected cells with specific JAK2 inhibitor, NSC 33994, confirmed that JAK2 regulates autophagy induction through JAK2-dependent pathway (Fig.6d). Therefore we tested the effect of NSC33994 on the viability of intracellular MTB in dithiopyridine and CI994-treated macrophages. Interestingly, inhibition of JAK2 signalling pathway by NSC33994 was found to be counteractive to the treatment with dithiopyridine/CI994 and led to the restoration of the viability of intracellular MTB. To investigate the role of STAT4, we knocked down *STAT4* in differentiated macrophages and infected them with virulent MTB. As expected, the MTB infected, *STAT4-*deficient cells did not exhibit the effect of dithiopyridine/CI994-induced MTB killing and confirmed the role of STAT4 in inducing IL-12-mediated autophagy (Fig. 6d).

**Figure 6.**
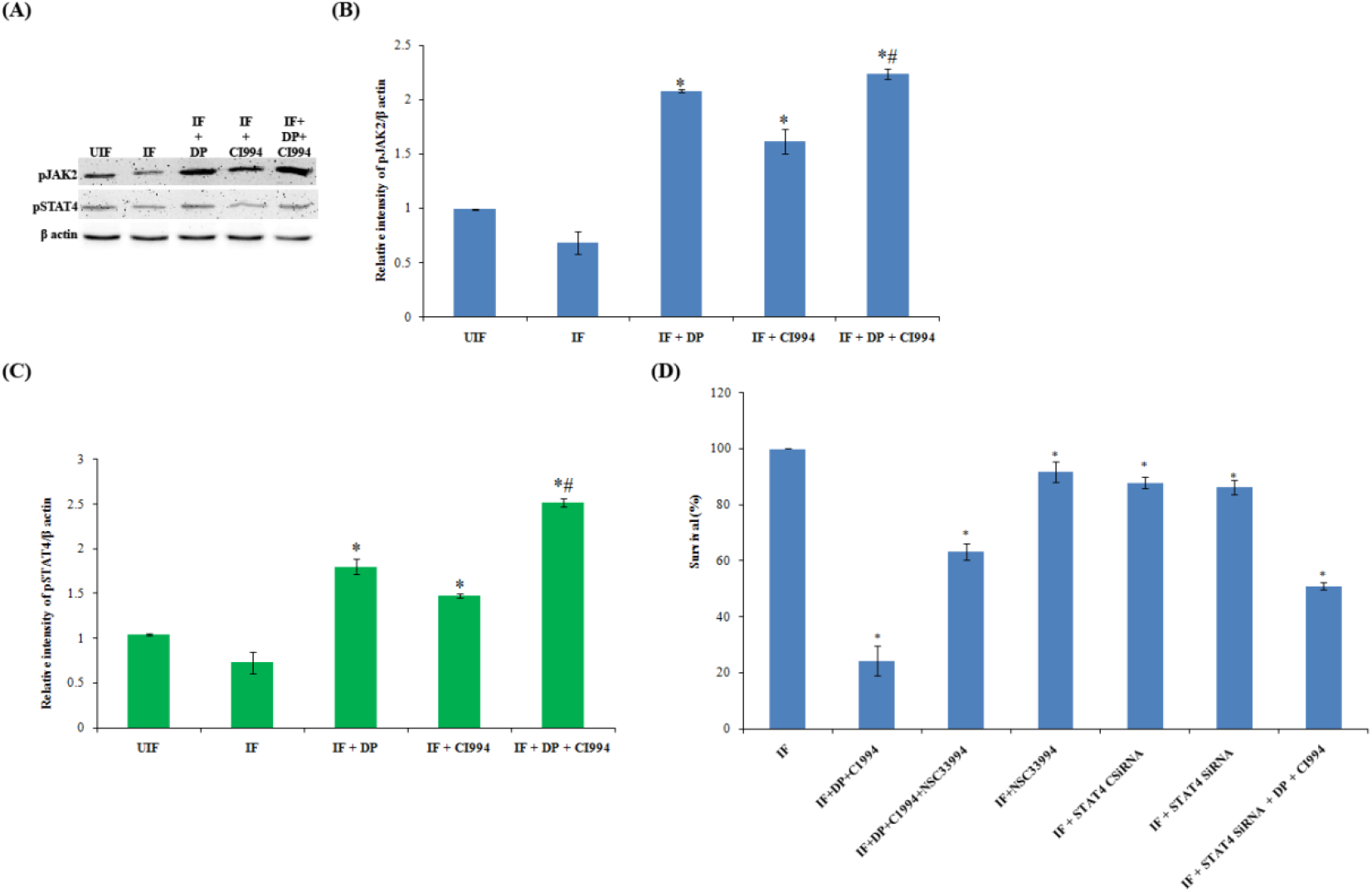
IL-12B mediated activation of autophagy depends on JAK-STAT pathway. (A) Status of pJAK2 and pSTAT4 in infected macrophages after treatment with dithiopyridine (DP) and CI994.Densitometric analysis of (B) pJAK2, and (C) STAT4 bands normalized with that of β-actin.Each value represents mean ± SD from triplicate measurements. *,#Band intensity is significantly different from infected (IF) and infected samples individually treated with inhibitors respectively, p≤0.05;(D) Infected THP1-derived macrophage cells were treated with JAK2 inhibitor NSC 33,994 along with DP and CI994. THP1-derived macrophages deficient inSTAT4 weretreated with DP and CI994. Intracellular bacterial viability was determined based on the number of CFUs. Isolation of bacilli from the macrophages was carried out at 24 h. The number of viable bacilli in each of the plate was assayed by plating lysed macrophages on 7H10 agar plates and incubating the plates at 37°C and counting the CFUs.Each value represents mean ± SD from triplicate measurements. * Survival is significantly different from infected (IF) sample, p≤0.05.

## Discussion

Earlier our laboratory had demonstrated that levels of HDAC1, a ubiquitous suppressor of gene expression, go up in macrophages infected with MTB, and it hypoacetylates histone H3 at the promoter of *IL-12B* gene (Chandran et al. 2015). Product of this gene plays a crucial role in initiating Th1 responses after infection with intracellular pathogens. HDAC1 is usually found in multi-protein corepressor complexes containing Sin3a, nucleosome-remodeling (NuRD) and CoREST, which are recruited to the promoter region of different genes by transcription factors like Sp1, Sp3, p53, NF-κB and YY1 (Delcuve et al, 2012). The SIN3A/HDAC complex is recognised as a global transcriptional corepressor. The Sin3A protein provides a platform for multiple protein interaction with its four paired amphipathic α-helix motif (Silverstein and Ekwall, 2005). HDAC1 forms a repressor complex and its function is regulated by the proteins it associates within the complex, which determine the enzymatic activity and recruitment to specific DNA sequences (Grignani et al, 1998). Phosphorylation of HDACs is the most widely studied post-translational modification, and the functional role of HDAC1, its cellular localization, and interaction with other proteins are regulated by this modification (Pflum et al, 2001). The identity and role of proteins associated with pHDAC1 during MTB infection and their role in the progress of infection and pathogenesis are not known. Time-course analysis revealed that HDAC1 phosphorylation is high at 24h of PI, and we presume that this modification contributes to its ability to bind to its associated proteins. In the present study, mass spectrometric analysis of immunoprecipitated HDAC1-interacting proteins revealed the presence of an unusual protein in the complex. We found that in addition to HDAC1 and Sinc3a, the repressor complex in MTB-infected macrophages contained the transcriptional repressor ZBTB25. We substantiated the association of ZBTB25 with HDAC1 and Sin3a by reciprocal co-immunoprecipitation and western blot analyses.

ZBTB25 is a transcriptional repressor, and the exact physiological function of ZBTB family is not well understood; a few ZBTB proteins have been reported to participate in tumour progression, chromatin remodeling, and in the development and differentiation of T and B cells (Chevrier et al, 2014; Lee & Maeda, 2012). It has been shown that transcriptional repressors participate in the progress of a variety of diseases (Xuan et al, 2013; Mao et al, 2013; He et al, 2016). In human lung adenocarcinoma epithelial cells infected with influenza virus ZBTB25 was found to interact with the viral RNA-dependent RNA polymerase (RdRp) and to enhance its transcription. In addition, ZBTB25 suppressed interferon production, further enhancing influenza viral replication (Chen et al, 2017). Nonetheless, the functional role of transcription repressors of ZBTB family involved in bacterial infection has not been described. ZBTB25 contains a DNA-binding zinc finger motif at the C-terminal, and a protein-binding BTB/POZ domain at the N-terminal. Proteins with these two conserved domains are categorized as POZ and Krüppel (POK) proteins, also called the Zinc finger and BTB domain (ZBTB) family (Krishna et al, 2003). Genomes of mouse and human code for more than 40 POK type proteins, which include promyelocytic leukaemia zinc finger proteins, ZBTB2, ZBTB7, ZBTB16 and ZBTB20 (Kelly et al, 2006; Costoya et al,2007; Kim et al; 2015; Wei et al, 2018).

The present study showed that ZBTB25 associated HDAC1 silencing complex is recruited to the *IL-12B* promoter and suppresses the expression of the *IL-12B* gene, thereby downregulating the host’s immune response. This generates a beneficial environment for MTB inside the macrophages. The protein binding domains of ZBTB25 interacts with HDAC1 and the DNA-binding domain binds to the *IL-12B* promoter sequences, respectively, since this protein has both DNA-binding and protein-binding domains (Krishna et al, 2003). We confirmed this interaction by in silico docking, immunoprecipitation and chromatin immunoprecipitation, and confocal microscopy. In addition we found that dithiopyridine, an inhibitor of ZBTB25, blocked the recruitment of HDAC1 silencing complex to *IL-12B* promoter. Dithiopyridine is a zinc ejector that inactivates the functionality of zinc finger domain (Lee et al, 2013), and has been used for the inactivation of the zinc-finger nucleocapsid of HIV-1 (Arthur et al, 1998). We further found that knocking down of ZBTB25 disrupts the recruitment of HDAC1 to the promoter of *IL-12B* promoter and subsequently derepressing the gene expression. Thus ZBTB25 plays a crucial role in the recruitment of HDAC1 silencing complex to *IL-12B* promoter in macrophages infected with MTB, in vitro. Because of its central role in the HDAC1 repression complex, we have also used a specific HDAC1 inhibitor CI994 (Beckers et al, 2007) in combination with dithiopyridine in infected macrophages. This combination treatment significantly reduced the number of viable MTB in infected macrophages. Phenylbutyrate (PBA), an HDAC inhibitor, has been shown to induce *CAMP/LL-37* gene expression in macrophages and inhibit MTB growth (Rekha et al, 2015).

The results from our study shows that the inhibition of ZBTB25 and HDAC1 increase the levels of IL-12p40 (a subunit of the cytokine IL-12) in MTB infected macrophages. Since IL-12 has been shown to be an activator of autophagy (Yang et al, 2018), we investigated if autophagy can be induced in THP-1-derived macrophages by treating them with dithiopyridine and CI994. Autophagy is one of the homeostatic machinery mediated through lysosomes and acts as an inherent defense mechanism against MTB infection. Host innate immunity uses autophagy to clear MTB and other intracellular pathogens (Shin et al, 2010; Rekha et al, 2015; Kim et al, 2019). Macrophages phagocytose and sequester MTB in phagosomes, which can then fuse with lysosomes for bacterial degradation. However, MTB has the ability to survive and grow inside macrophages by preventing phagosome-lysosome fusion (Gutierrez et al, 2004). Autophagy induction can be harnessed toward stimulating macrophage defense against MTB (Xu et al, 2007). Gutierrez *et al*. (2004) showed that starvation or treatment with rapamycin increases MTB colocalization with LC3 and beclin-1, and transfers MTB to phagolysosomes for subsequent autophagy. It has been shown that anti-protozoal drug nitazoxanide and its active metabolite tizoxanide strongly stimulate autophagy and inhibit signaling by mTORC1, a major negative regulator of autophagy and inhibit MTB proliferation more potently in infected human THP-1 cells and peripheral monocytes (Lam et al, 2012). Anti-mycobacterial drugs like isoniazid and pyrazinamide have been shown to induce autophagy and phagosomal maturation in MTB-infected host cells (Kim et al, 2012). Loperamide and cholesterol-lowering drug statin were also reported to induce autophagy and phagosomal maturation and reduce intracellular growth of MTB in infected macrophages, suggesting that host autophagy plays a key role in host immune responses during drug administration against TB (Juárezet al, 2016, Parihar et al, 2014). We here showed that treatment of THP-1-derived macrophages with dithiopyridine and CI994 restores the levels of IL-12p40 and leads to the activation of major autophagic markers, enhances intracellular LC3 distribution, and causes colocalization of MTB with LC3. These observations suggest that treatment of macrophage with dithiopyridine and CI994 could be an effective antimicrobial strategy against intracellular MTB by inducing autophagy. Interestingly Jung et al. (2014) showed that IL-12 is involved in antimicrobial action by activating the fusion of phagosome and lysosome via IFN-γ. Although IL-12-induced autophagy has been reported in breast cancer cells, human macrophages, and lung epithelial cells (Lin et al, 2017; Yang et al, 2018), the precise function of IL-12 in inducing this process is not yet clearly understood.

IL-12 activates tyrosine phosphorylation of janus family tyrosine kinases JAK2 and Tyk2, suggesting the involvement of these kinases in the biochemical response to IL-12. The substrates for the JAKs are the family of latent cytoplasmic transcription factors termed STATs (Bacon et al, 1995). The presence of the phosphotyrosine dimer in STATs is vital in target gene activation (Bromberg, 2001). To study the mechanism by which IL-12p40 activates autophagy, we evaluated the status of JAK2/STAT4 in the MTB infected macrophages. Enhanced phosphorylation of JAK2/STAT4 was observed in infected macrophages upon dithiopyridine/CI994 treatment. JAK2 inhibition by specific inhibitor NSC33994 and knocking down of *STAT4* also led to an increase in intracellular survival of MTB through inhibition of dithiopyridine/CI994-induced autophagy. Taken together these results clearly demonstrate that dithiopyridine and CI994-mediated restoration of IL-12p40 levels induces autophagy by phosphorylating JAK2 and STAT4 that eventually kills intracellular MTB.

Host-directed therapy (HDT) is an emerging treatment method for TB, where host response is enhanced by treatment with different drugs, with or without antimicrobials, to achieve better management of TB. HDT drugs targets autophagy, vitamin D pathway, and anti-inflammatory responses (Kolloli et al, 2017). Jayaswal et al (2010) highlighted the importance of host factors that regulate intracellular survival of MTB through an SiRNA screen in J774.1 murine macrophage cells. They could identify host factors such as TGFβRI and CSNK1, and their inhibition substantially reduced the number of intracellular MTB in J774.1 cells as well as in primary murine macrophages. Pharmacological inhibition of the AKT/mTOR pathway in host reduces intracellular growth of MTB both in vitro in human PBMCs, and *in vivo* in a model of murine tuberculosis (Lachmandas et al, 2016). Targeted inhibition of NFκB activation decreases viability of intracellular MTB in human macrophages (Bai et al, 2013). HDAC6 inhibitor Tubustatin A has been shown to reduce MTB survival in an *in vivo* mouse model (Wang et al, 2018). The pan-HDAC inhibitor trichostatin A and class IIa HDAC inhibitor TMP195 have been shown to be very effective in reducing MTB infection in primary human macrophages, and *Mycobacterium marinum* infection in zebrafish model (Moreira et al, 2020). Rao et al (2018) evaluated the possibility of the histone deacetylase inhibitors valproic acid (VPA) and suberoylanilide hydroxamic acid (SAHA) to enhance the activity of first-line anti-TB drugs to inhibit the growth of intracellular MTB. Thus there has been considerable interest in using inhibitors of HDACs (HDACi) for TB treatment. However, considering the enormous number of genes that are regulated by different HDAC enzymes, the utility of HDAC inhibition as a safe therapeutic strategy presents numerous challenges. This necessitates identification of disease-specific host targets to treat infectious diseases. In this context, our discovery that transcriptional repressor protein ZBTB25 is directly associated with HDAC1 repressor complex in MTB-infected macrophages, and its inhibition or knocking down significantly reduces the number of intracellular MTB, makes this protein an interesting candidate for possible therapeutic intervention. More studies with infected animal models are required to provide an answer to the question of whether inhibition of ZBTB25 would be a better option than inhibiting HDACs to control MTB infection in humans as a host-directed therapeutic strategy.

## Materials and Methods

### Bacterial strains

MTB strains H37Rv (virulent) and MTB H37Rv expressing GFP constitutively were used in the current study, and these MTB strains were handled in a Biosafety Level 3 (BSL3) facility. Middlebrook 7H9 broth (Difco) added with 10% OADC (Becton Dickinson), 0.4% glycerol, and 0.05% Tween 80 were used to grow the MTB strains. Bacteria in the mid-log phase were centrifuged, washed, and resuspended in RPMI medium containing FBS (10%). The suspension was dispersed by aspirating several times and vortexed until no bacterial clumps were visible, and allowed to stand for 5 min. The upper half of the suspension was collected and quantified (absorbance of 0.15 corresponds to ~3 × 10^8^ bacteria at 600 nm) to perform various experiments.

#### Resazurin microtitre assay (REMA)

REMA was performed as described by Martin *et al.* (2003). In short, the turbidity of MTB cultures was adjusted to 0.2 units at 600 nm which correspond to 3 × 10^8^ bacteria/mL and was diluted (20 times). The diluted bacterial cells (100 μL) were added to the 96 well microtitre plate (Nunclon, Thermo Fisher Scientific, Denmark) containing 100 μL of dithiopyridine, CI994 or rifampicin solutions. The microtitreplates were incubated for 24 h for 7 days with appropriate controls. After incubation, minimum inhibitory concentration (MIC) was determined by adding resazurin (30 μL of 0.02% aqueous solution). Blue colour of the culture would indicate inhibition of bacterial growth whereas pink colour would indicate viable bacteria.

### Cytotoxicity assay by MTT

The MTT assay was performed as described by Mosmann (1983). Briefly, 10 000 cells were seeded per well in a 96 well plate. They were treated with dithiopyridine and CI994 and incubated for 48 h at 37 °C in the presence of 5% CO_2_. Ten microlitre of MTT (3-(4,5-dimethylthiazole-2-yl)-2,5-diphenyl tetrazolium bromide) dye (5 mg mL−1) was added to each well and after 4 h, the reaction was terminated by adding 200 μL of DMSO and this dissolves the formazan crystals. This was read at 540 nm on an ELISA reader (Bio-Rad).

### Infection of THP-1-derived macrophage cells with *M. tuberculosis* H37Rv

RPMI 1640 medium containing FBS (10%), L-glutamine (2 mM), HEPES (25 mM) and Na_2_CO_3_ (1.5 g/L) was used to culture THP-1 cells at 37 °C and CO_2_ (5%). Upon treatment of THP1-derived macrophages with phorbitol 12-myristate 13-acetate (PMA) (20 ng/mL) for 24 h they get differentiated into macrophages. The differentiated cells were mixed with MTB at 20:1 multiplicity of infection (MOI) and were incubated for 4 h at 37 °C in the presence of 5% CO_2_ to facilitate phagocytosis. Cells were then washed four times with complete RPMI containing gentamycin (10 μg/mL) to remove extracellular bacilli and fresh medium was added to the cells. For experiments in which inhibitors were used, specific inhibitors were added after phagocytosis. Final concentrations of the inhibitors were: dithiopyridine, 20 μM/ml; CI994, 15 μM/ml; Rapamycin, 100 nM/ml.

Apart from THP-1-derived macrophage cells, primary macrophages obtained from peripheral blood mononuclear cells (PBMC, Sigma) were also used in the study. PBMCs were incubated at 37 °C in 5% CO_2_ for 2 h. The non-attached cells were removed by washing thrice with RPMI-1640. The remaining attached cells were grown in RPMI supplemented with FBS (10%), incubated for 4 days to differentiate into macrophages, and further infected with MTB.

### Si RNA transfection of macrophages and MTB infection

THP-1 cells grown in Opti-MEM medium (Invitrogen, Carlsbad, USA) were differentiated and transfected (Hiperfect transfection reagent (Qiagen, Valencia, USA) with 60 pmol of siRNA (Santa Cruz Biotechnologies, USA) for 36 h. Transfected cells were infected with MTB H37Rv at an MOI of 20:1. Cells transfected with scrambled siRNA served as control.

### Confocal microscopy

Differentiated THP-1 cells were infected with MTB H37Rv and MTB GFP-H37Rv, and incubated for 4 h for phagocytosis. The extracellular bacilli were removed, cells were fixed with parafolmadehyde (4%), incubated with methanol for 20 min at −20 °C, and then blocked with PBS [containing BSA (3% w/v), and Triton X-100 (0.3% v/v)]. The cells were incubated for 1 h at room temperature with anti-rabbit or anti-mouse primary antibodies against pHDAC1, ZBTB25, LC3, ATG5 and BECN1, washed, and incubated with FITC-conjugated anti-rabbit or anti-mouse secondary antibody (Sigma-Aldrich) or Alexa-488/Alexa-592 anti-rabbit or anti-mouse secondary antibody (Thermo Fischer, US) for 1 h at room temperature. Cells were counterstained with DAPI for DNA. To detect lysosomes, cells were incubated with LysoTracker-Red (Thermo Fisher Scientific). Rapamycin (Sigma Aldrich) was used as a positive control for autophagy induction.

### Immunoprecipitation (IP)

Differentiated THP-1 cells (infected and uninfected) were harvested, washed with PBS, and then extracted with IP lysis buffer for 30 min at 4 °C. Specific primary antibodies or rabbit or mouse secondary immunoglobulin G (IgG) was added to MTB infected and uninfected cell lysate and incubated overnight at 4 °C. Then Protein-A/G coated sepharose beads were added to the antibody-protein mixture for 2 h at 4 °C. Beads were washed four to five times with lysis buffer and eluted in 0.2 M glycine and precipitated with TCA/acetone and dissolved in 50 mM ammonium carbonate and subjected to LC-MS/MS. For western blot the immuncomplexes were eluted in SDS/PAGE sample buffer (0.0005% Bromophenol blue, 10% Glycerol, 2% SDS, 63 mM Tris-HCl pH 6.8).

### Chromatin immunoprecipitation (ChIP)

ChIP was carried out based on the manufacturer’s protocol (Abcam, UK). In brief, the infected and uninfected macrophages were treated with 4% formaldehyde in PBS for 10 min and washed with PBS. Cells were lysed and the chromatin was sheared by sonication (Bioruptor, Diagenode, Belgium) with pulse conditions 30 S on, 30 S off for 25 cycles at 4 °C followed by centrifugation (14000 rpm × 15 min at 4 °C). Antibodies against HDAC1, ZBTB25, Sin3a, or IgG were used to immunoprecipitate the chromatin fragments in the supernatant by overnight incubation using protein A sepharose beads (Abcam, UK) at 4 °C.The beads were washed twice with wash buffer, DNA was eluted off the beads, and the purified DNA was subjected to PCR. Oligonucleotide primers used for PCR amplification of specific gene promoters are listed in Table 2 in Supplementary Material.

### Molecular Docking

To infer the interaction between HDAC1 and ZBTB25, we performed docking studies by utilizing Autodock 4 software (Morris et al, 2009).

### Western blotting for specific molecules

Nuclear extracts (using NE-PER™ nuclear Extraction Kit (Thermo Fischer Scientific, USA) and whole-cell lysates (using RIPA buffer, Sigma) were prepared from MTB-infected macrophages (MOI 20:1). Proteins were separated on SDS-polyacrylamide (12%) gels, transferred to Immobilon-P polyvinylidenedifluoride membranes (Millipore, Billerica, MA), and probed with protein-specific primary antibody. Then anti-rabbit or anti-mouse HRP-conjugated secondary antibody was added, and the target proteins were visualized by chemiluminescence, and image quantification was carried out using ImageJ software (NIH, USA). Relative intensity of specific proteins was plotted in correspondence to band intensity of Histone-H3 or actin.

### Quantitative real-time PCR (qPCR)

The total RNA was isolated from macrophages (infected and uninfected) using Nucleo spin RNA extraction kit (Macherey-Nagel, Germany) according to the manufacturer’s protocol. RNA (1 μg) was converted to cDNA using the GoScript™ Reverse Transcription System (Promega, Madison, WI). The cDNA was synthesised was used for qPCR using the iQTM SYBR® Green Supermix (Bio-Rad Laboratories, Hercules, CA) on an iCycleriQ™ Real-Time PCR Detection System (Bio-Rad). Fold changes in gene expression levels were calculated and were normalized to human actin. The list of qPCR primers (from Sigma-Aldrich, USA) is shown in Table 2 in Supplementary Material.

### ELISA

The IL-12p40 in cell supernatants of infected and uninfected macrophages were quantified using BD OpteEIA™ Human IL-12(p40) ELISA kit (Abcam, UK) as per the manufacturer’s directives.

### Intracellular survival assays for MTB

The THP-1-derived macrophage cells were infected with MTB as described elsewhere. After 4 h, cells were washed four times with complete RPMI containing gentamycin (10 μg/mL) to remove extracellular bacilli and fresh medium was added. Cells were harvested at 24 h after infection. The monolayers of macrophages were washed with PBS, lysed with SDS (0.06%) in 7H9 medium, and the lysate was spread on 7H10 agar plates and incubated at 37 °C to determine the intracellular colony forming units (CFU). After 3 weeks, the bacterial colonies on the plates were counted. The experiment was performed in triplicates.

### Statistical analysis

The data were expressed as mean values with their standard deviations. The results were analyzed by non-parametric analysis of variance (ANOVA). A p < 0.05 was considered significant.

## Acknowledgment

Aravind Madhavan would like to acknowledge the Department of Health Research, Government of India, Science Engineering and Research Board (Department of Science and Technology, Government of India) and Department of Biotechnology, Government of India for post-doctoral fellowships. Arun KB would like to acknowledge the Department of Health Research, Government of India, Science Engineering and Research Board (Department of Science and Technology, Government of India), and Kerala Biotechnology Commission (Kerala State Council for Science and Technology, Government of Kerala) for post-doctoral fellowships. R.A.K is grateful to the Department of Biotechnology, Government of India for financial support. The authors thank the staff of confocal microscopy facility, RGCB, for their support in imaging. The authors also thank Sivakumar K.C for the support in molecular docking studies.

---

**Supplementary Table 1:** Identification of host proteins that interact with HDAC1 in macrophages infected with MTB by LC-MS/MS

| **Protein Name** | **Molecular Weight** | **Uniport ID** | **Number of Peptides from LC MS/MS** |
| --- | --- | --- | --- |
| Histone deacetylase 1 | 55.103 | Q13547 | 14 |
| Histone deacetylase 2 | 55.364 | Q92769 | 10 |
| Zinc finger and BTB domain-containing protein 25 | 48.990 | P24278 | 12 |
| Paired amphipathic helix protein Sin3a | 145.175 | Q96ST3 | 11 |

**Supplementary Table 2:**
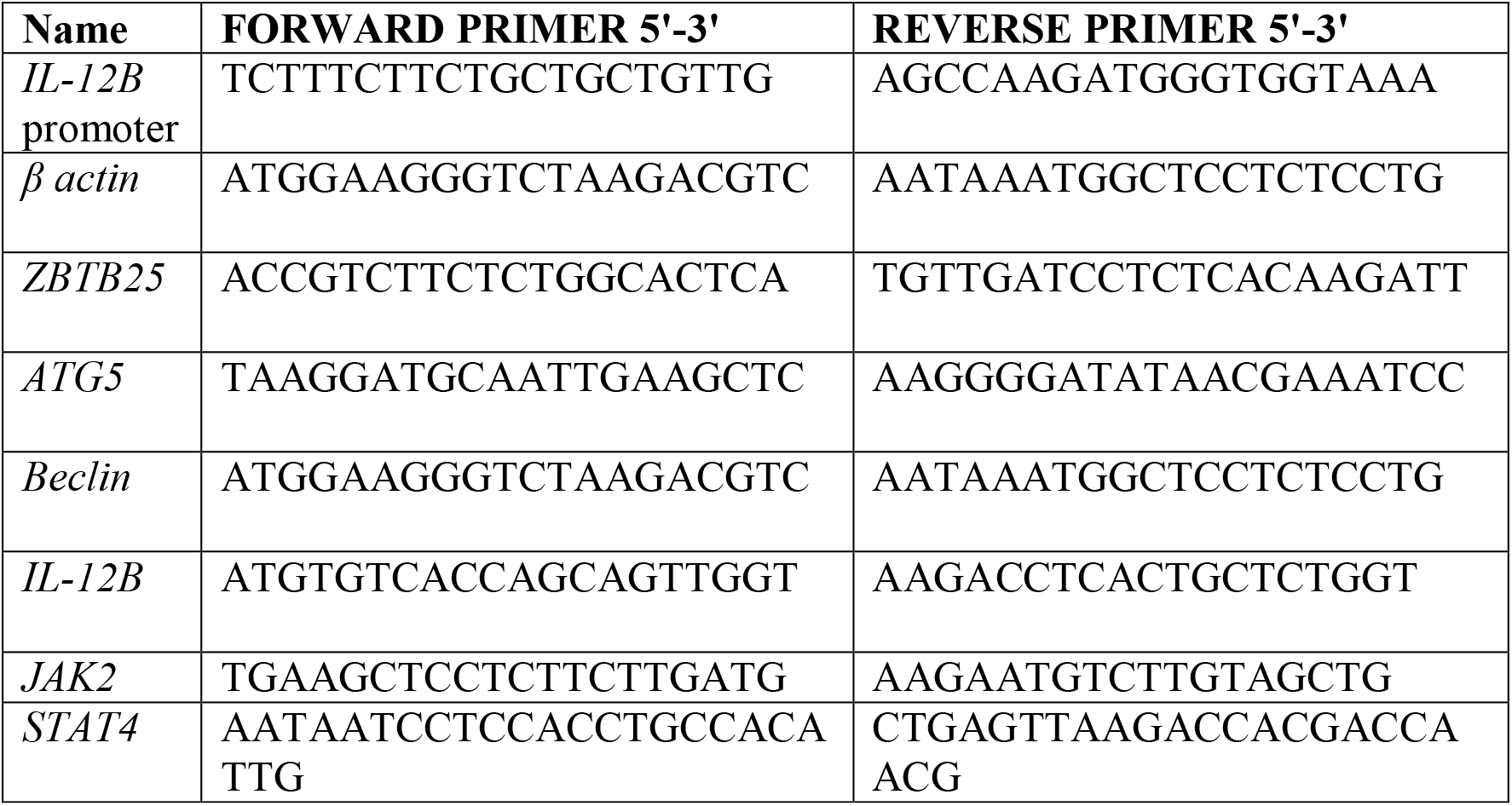
List of oligonucleotide primers used in this study

**Supplementary Figure 1:**
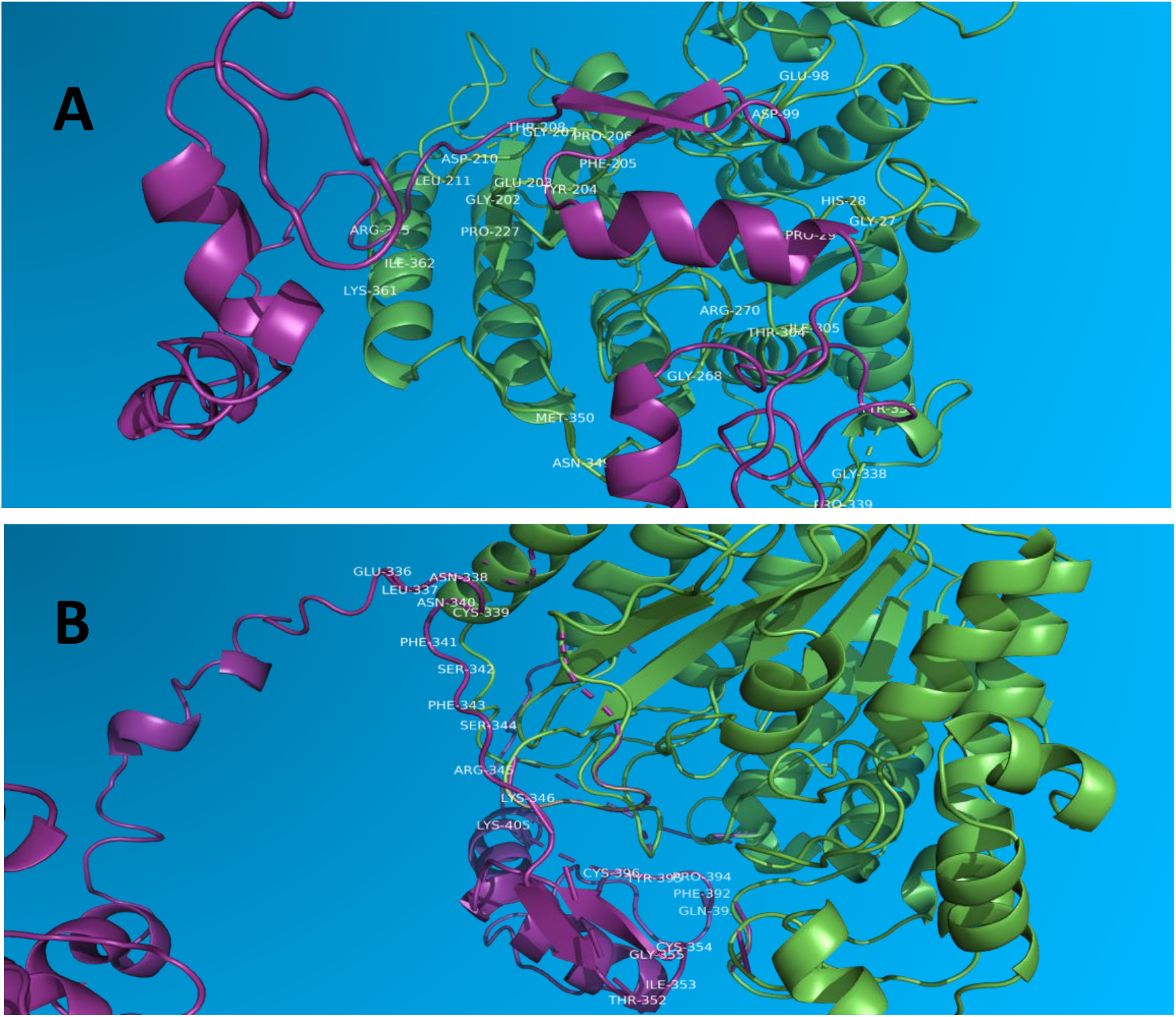
Docking analysis shows the interacting residues of (A) HDAC1(green) and (B) ZBTB25 (Purple) using Autodock 4.2.

**Supplementary Figure 2:**
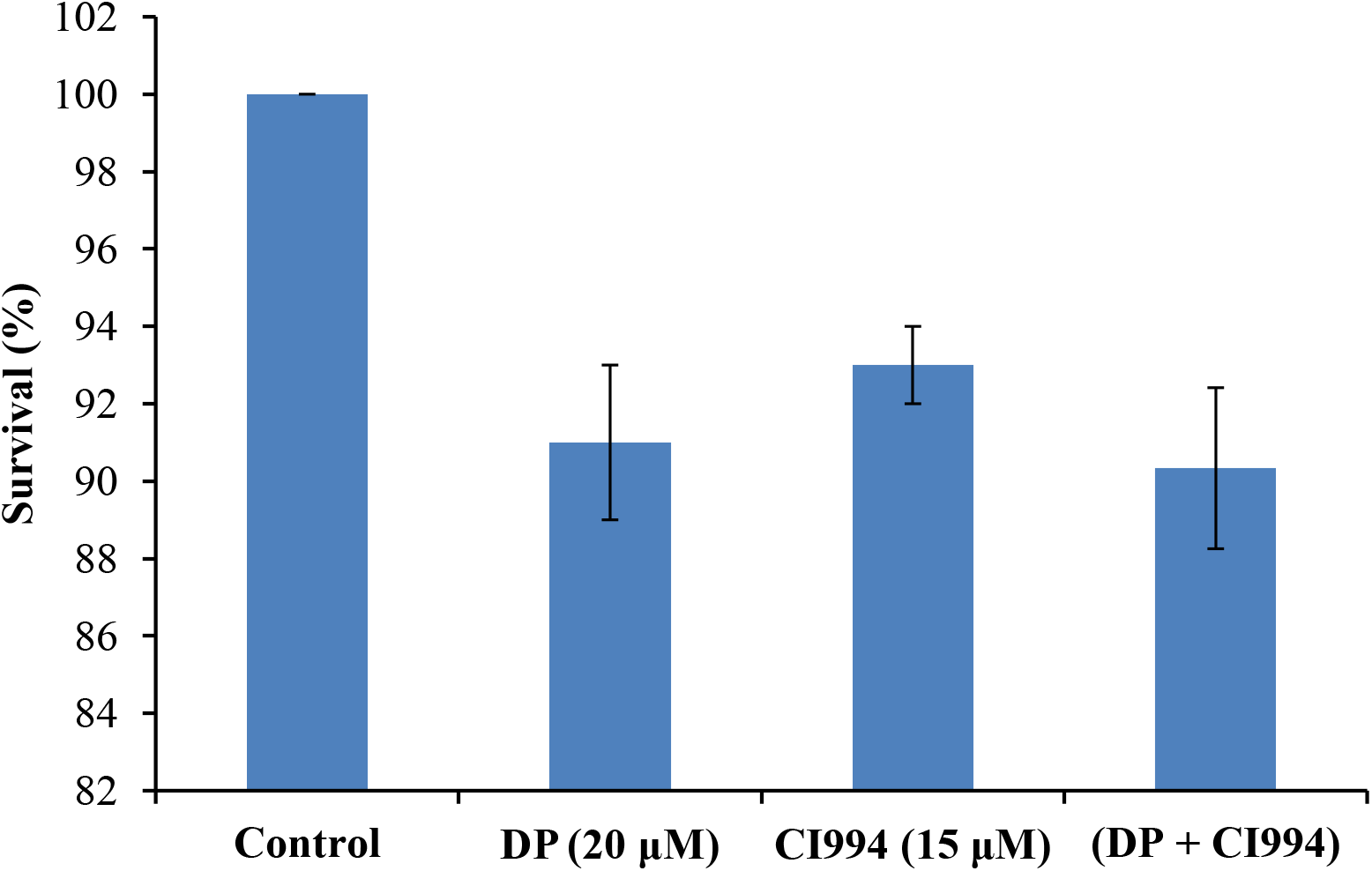
MTT assay: viability of THP1-derived macrophages upon treatment with dithiopyridine (DP) and CI994 at 48 h.

**Supplementary Figure 3.**
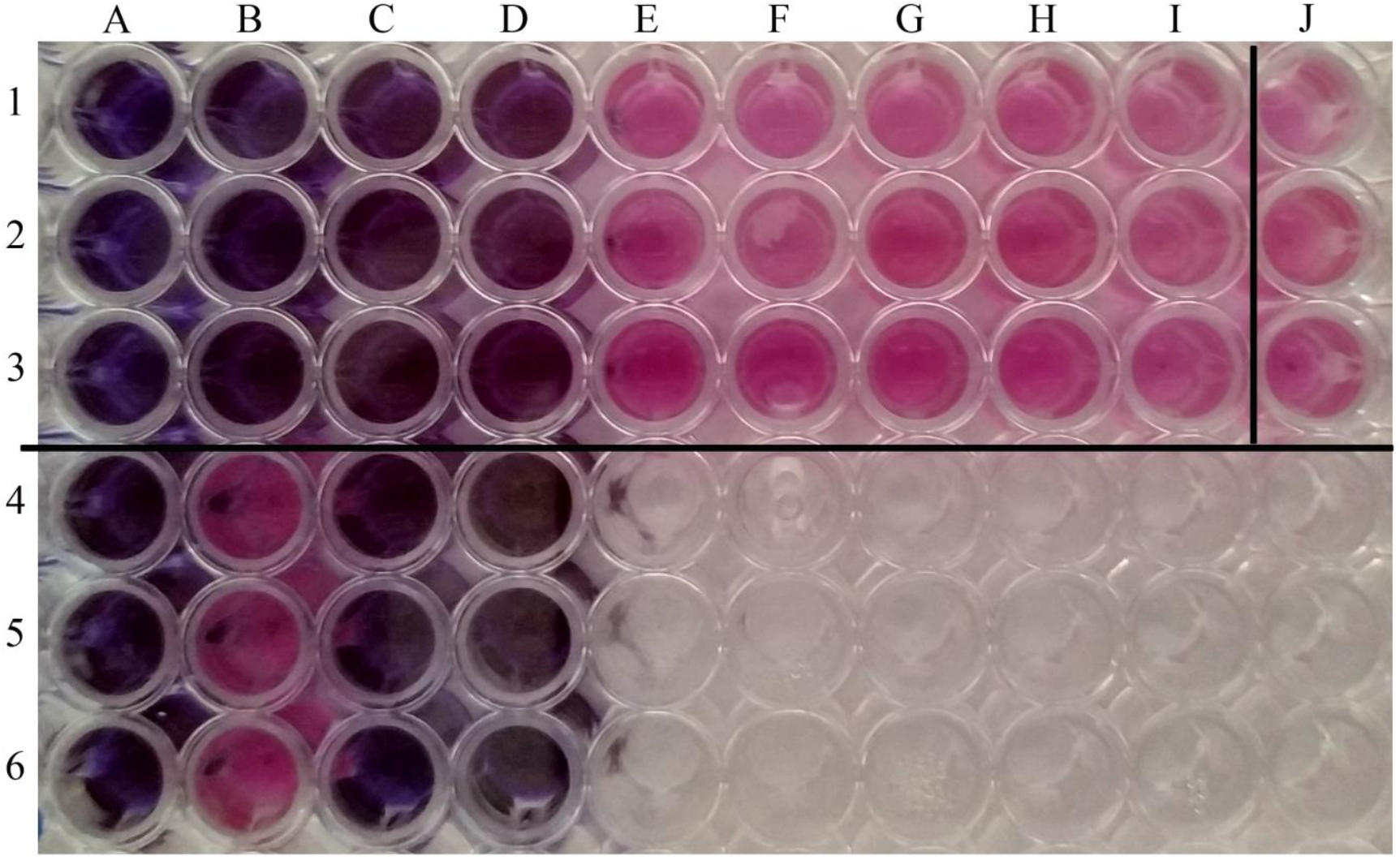
Resazurinmicrotiter assay(REMA) to determine the minimum inhibitory concentration(MIC) of DP against MTB. Wells 1A to 1I contain *M. tuberculosis* H37Rv treated with decreasing concentrations of DP (1000μM-3.90μM).Wells 2A to 3I represent duplicates of the same. Wells 1J,2J &3J contain DMSO control. Wells 4A, 5A & 6A contain media alone. Wells 4B,5B &6B contain media with bacteria.Wells 4C, 5C,6C contain media with drug alone. Wells 4D,5D & 6D contain bacteria treated with rifampicin (1 μg/mL).

**Supplementary Figure 4.**
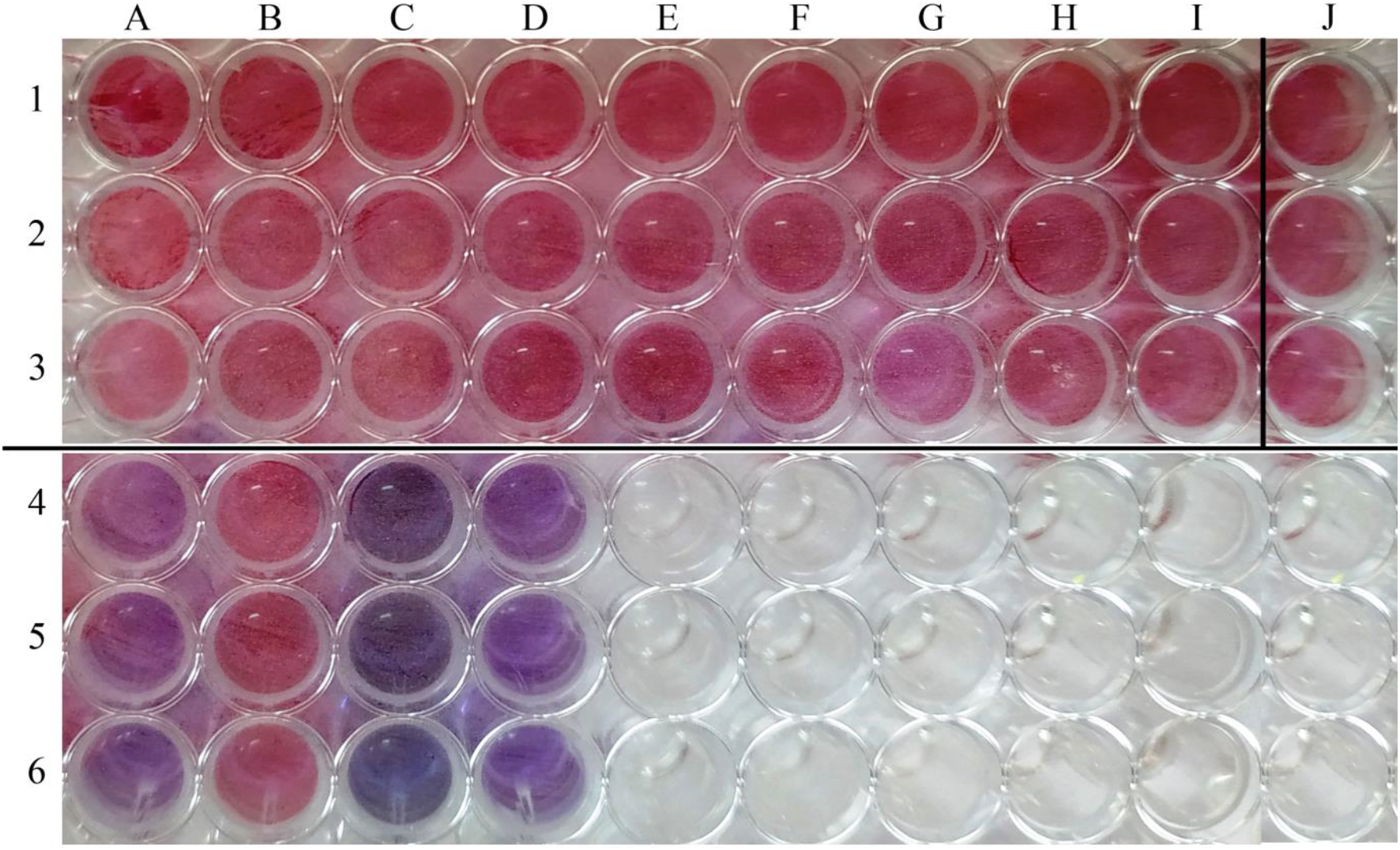
REMA of MTBin the presence of CI994. Wells 1A to 1I contain *M. tuberculosis* H37Rv treated with decreasing concentrations of CI994 (1000μM-3.90μM).Wells 2A to 3I represent duplicates of the same. Wells 1J,2J &3J contain DMSO control. Wells 4A, 5A & 6A contain media alone. Wells 4B,5B,6B contain media with bacteria. Wells 4D,5D & 6D contain bacteria treated with rifampicin (1 μg/mL). Wells 4C, 5C, 6C contain media with drug alone.

**Supplementary Figure 5.**
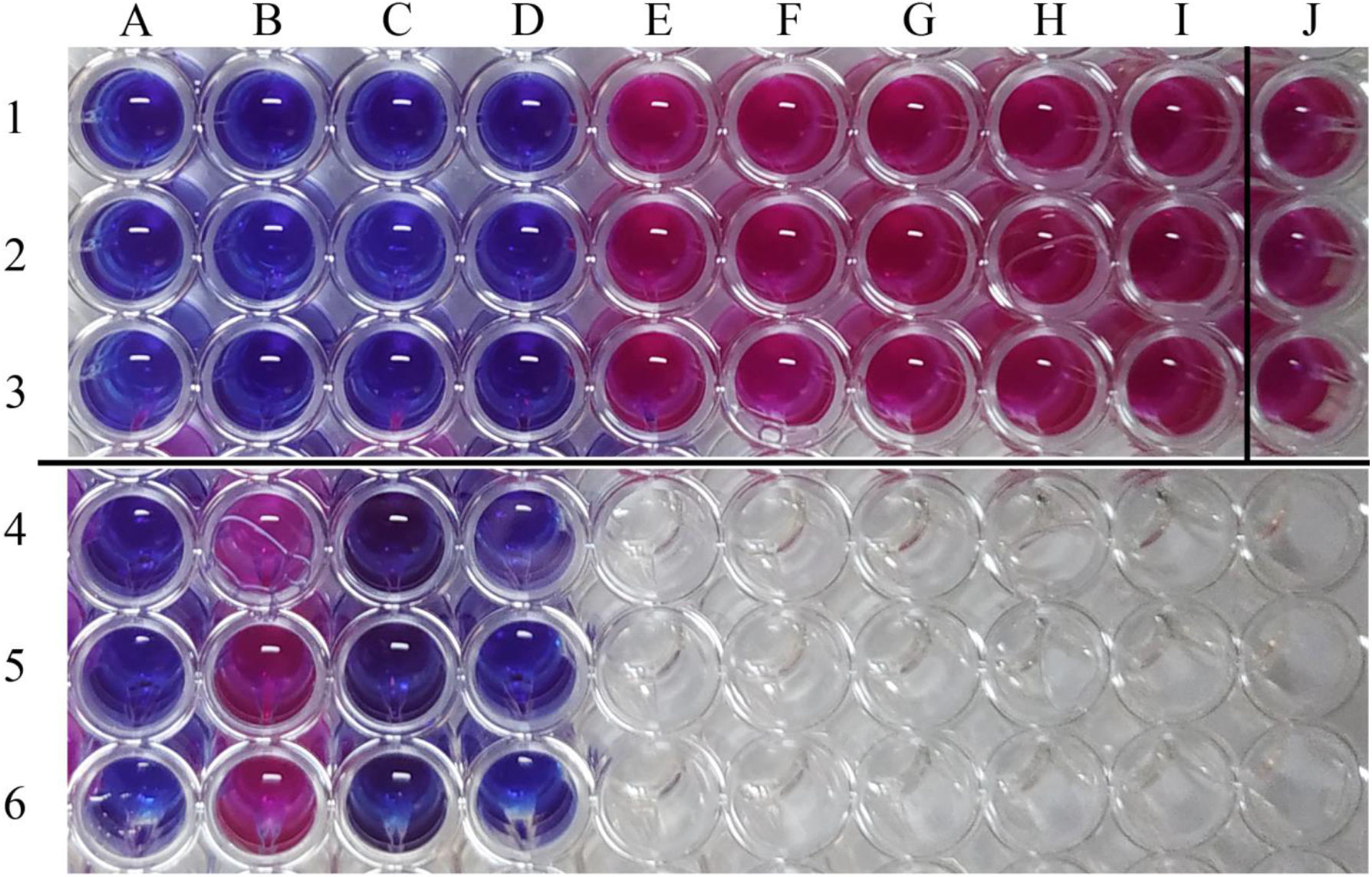
REMA of MTBin the presence of DP and CI994. Wells 1A to 1I contain *M. tuberculosis* H37Rv treated with decreasing concentrations of DP and CI994 (1000μM-3.90μM).Wells 2A to 3I represent duplicates of the same. Wells 1J,2J &3J contain DMSO control. Wells 4A, 5A & 6A contain media alone. Wells 4B,5B,6B contain media with bacteria. Wells 4D,5D & 6D contain bacteria treated with rifampicin (1 μg/mL). Wells 4C, 5C,6C contains media with drug alone.

**Supplementary Figure 6:**
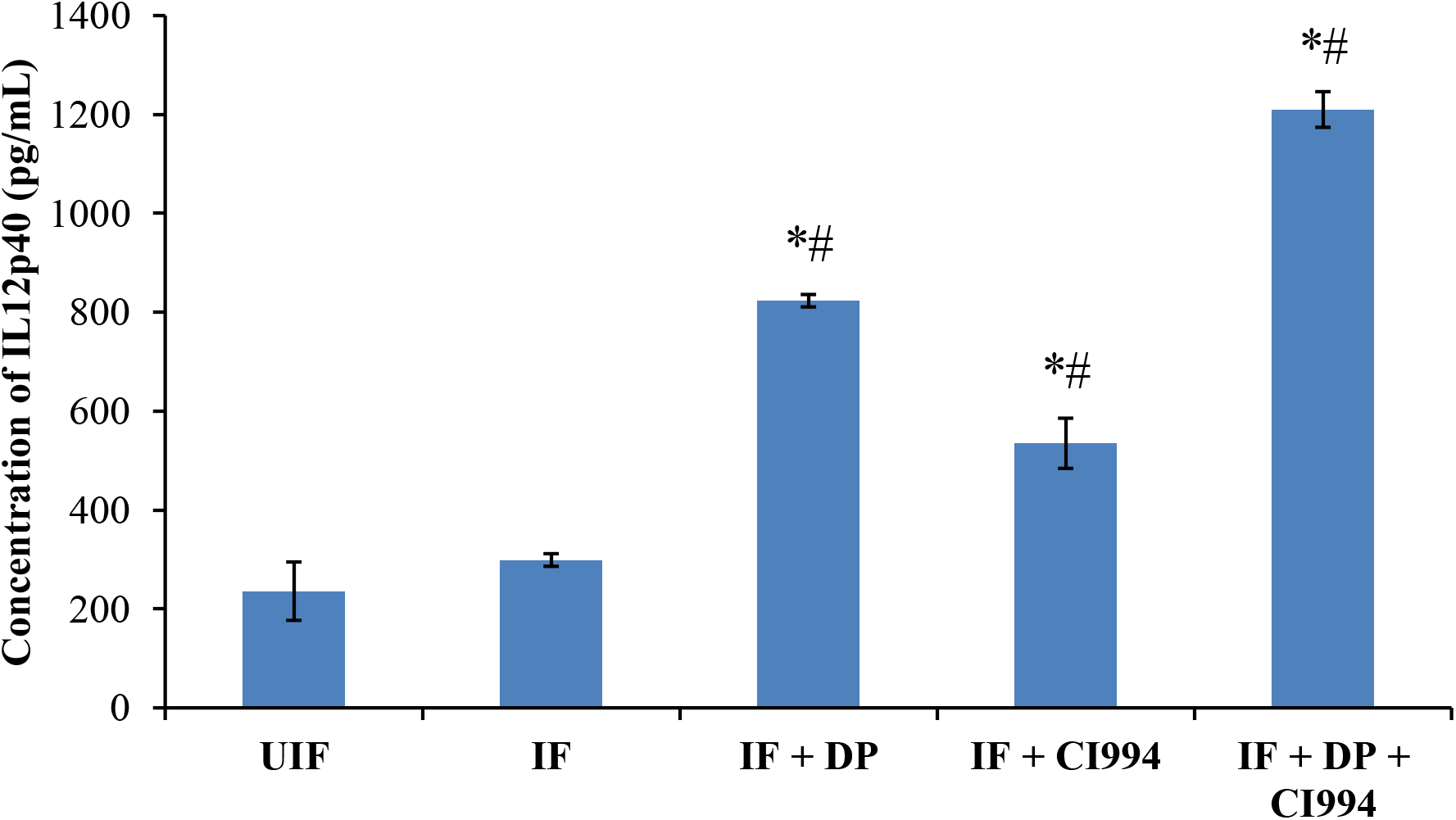
ELISA of IL-12p40 in PBMC after the infected cells were treated with DP and CI994.

**Supplementary Figure 7:**
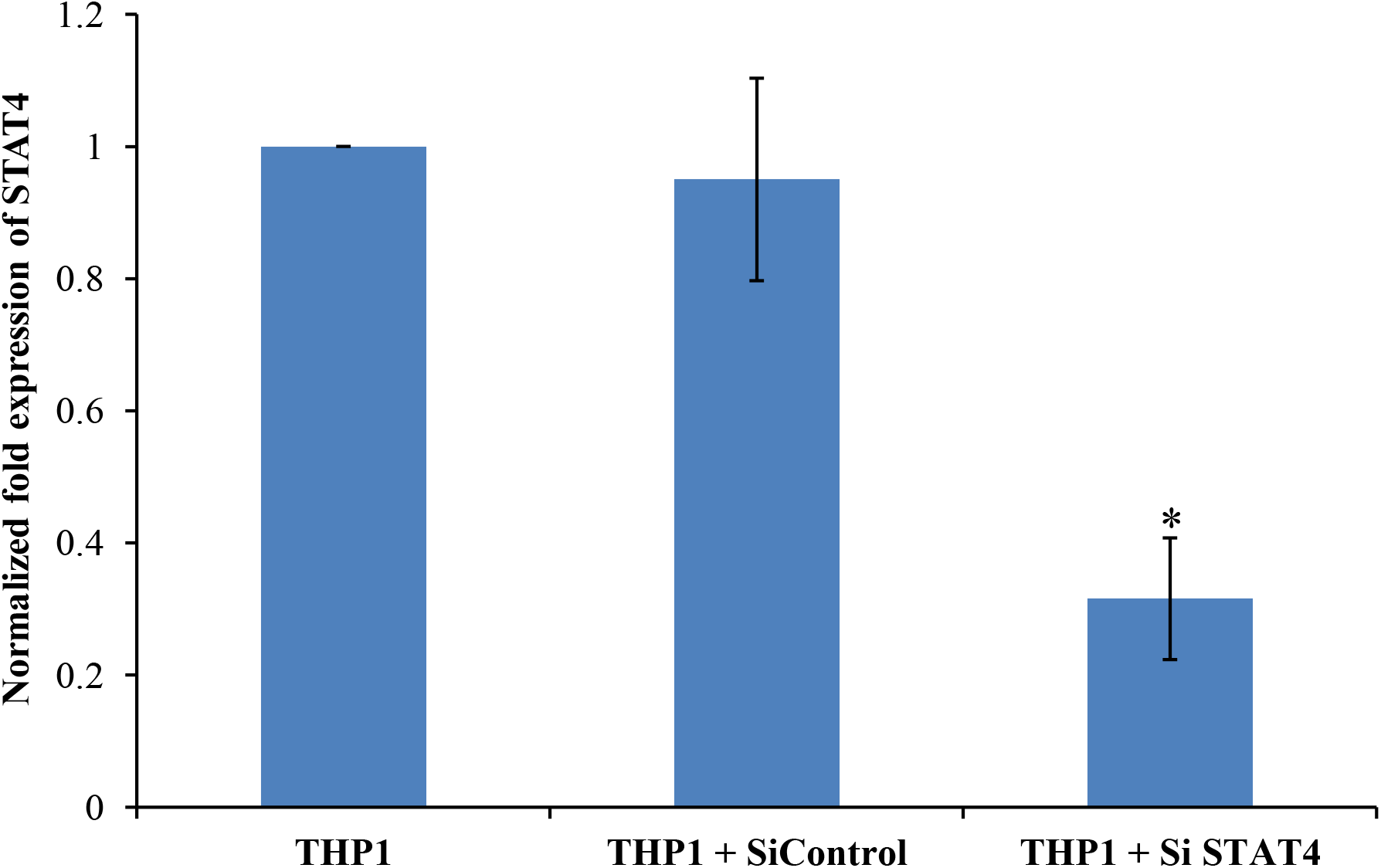
Efficiency ofknock down was confirmed by qPCR. THP1-derived macrophage cells were transfected with *STAT4*SiRNA or scrambled SiRNA control using Hiperfect transfection reagent.

